# Ubiquitin specific protease 1 expression and function in T cell immunity

**DOI:** 10.1101/2021.04.08.439046

**Authors:** Kyla D Omilusik, Marija S Nadjsombati, Tomomi M Yoshida, Laura A Shaw, John Goulding, Ananda W Goldrath

## Abstract

T cells are essential mediators of the immune responses against infectious diseases and provide long-lived protection from reinfection. The differentiation of naive T cells to effector T cells and subsequent differentiation and persistence of memory T cell populations in response to infection is a highly regulated process. E protein transcription factors and their inhibitors, Id proteins, are important regulators of both CD4^+^ and CD8^+^ T cell responses; however, their regulation at the protein level has not been explored. Recently, the deubiquitinase USP1 was shown to stabilize Id2 and modulate cellular differentiation in osteosarcomas. Here, we investigated a role for Usp1 in posttranslational control of Id2 and Id3 in T cells. We show that Usp1 was upregulated in T cells following activation *in vitro* or following infection *in vivo*, and the extent of Usp1 expression correlated with the degree of T cell expansion. Usp1 directly interacted with Id2 and Id3 following T cell activation. However, Usp1-deficiency did not impact Id protein abundance in effector T cells or alter effector CD8^+^ T cell expansion or differentiation following a primary infection. Usp1 deficiency did result in a gradual loss of memory cells over time and impaired accumulation and altered differentiation following a secondary infection. Together, these results identify Usp1 as a player in modulating recall responses at the protein level and highlight differences in regulation of T cell responses between primary and subsequent infection encounters. Finally, our observations reveal that differential regulation of Id2/3 proteins between immune vs non-immune cell types.

## Introduction

T cells are key components of the adaptive immune response to combat infectious disease. In the event of infection, naive T cells are induced to proliferate and differentiate into armed effector cells that can directly eliminate pathogen as well as enhance and direct the function of other immune cells. Upon the resolution of infection, a fraction of the antigen-specific cells persist indefinitely as memory T cells, providing protection from reinfection. Functional and phenotypic heterogeneity exists within both the CD4^+^ and CD8^+^ effector T cell populations. Naive CD4^+^ T cells can differentiate into different T helper lineages, including Th1, Th2, Th17, T regulatory and T follicular helper (Tfh) lineages depending on the context of the infection or inflammation (1). Two major subsets of effector CD8^+^ T cells can be identified by differential expression of CD127 and KLRG1, where KLRG1^hi^CD127^lo^ define terminal effector (TE) and KLRG1^lo^CD127^hi^ delineate memory precursor (MP) cells (2). Both populations undergo contraction as the infection is cleared; however, the TE subset continues to contract over the months following antigen exposure while the MP subset provides stable, persistent memory (3, 4). The long-lived circulating memory T cell pool has also been broadly classified into 2 main subsets: effector memory (T_EM_) and central memory (T_CM_) T cells (5, 6). T_EM,_ considered to be shorter-lived and more terminally differentiated, display enhanced effector-like properties and circulate through the vasculature and tissue while T_CM_ exhibit heightened homeostatic proliferation, multipotency, longevity and recall potential and mediate their localization to lymph nodes via expression of CD62L and CCR7 (5-8). Within the peripheral CD8^+^ T cell memory pool, a population of long-lived cells that are terminally differentiated but with potent effector functions, t-T_EM,_ have also been described (9, 10)

Heterogeneous memory T cell fates are directed and maintained by distinct transcriptional programs that are induced in activated T cells (11-15). E box-binding transcription factors and their inhibitors, Id proteins, are essential regulators of both CD4^+^ and CD8^+^ T cell differentiation (16-23). Increased E protein activity in CD8^+^ T cell shortly after activation supports a transcriptional program that promotes MP cell formation and E protein-deficient CD8^+^ T cells favor production of KLRG1^hi^ TE and effector-memory cells (17). Id2 and Id3 inhibit E protein activity to modulate T cell differentiation in seemingly opposing fashion. Effector CD8^+^ T cells require Id2 to accumulate and persist long term, and Id2-deficiency results in almost exclusive formation of the MP subset (16, 18-20). Conversely, Id3-expressing effector CD8^+^ T cells exhibit a similar transcriptional gene-expression profile to long-lived memory cells, preferentially differentiate into memory cells, survive longer, and responded better to secondary challenge compared to effector cells that do not upregulate Id3 (22).Within the CD4^+^ T cell helper and memory populations, E and Id proteins are critical for T helper cell specification. E protein deficiency promotes enhanced Treg cell development while Id2 and Id3 deletion impair differentiation and localization of Foxp3^+^ Treg cells and loss of Id2 results in reduced survival and function of adipose tissue Treg cells (24-27). Furthermore, CD4^+^ T cells lacking Id2 or Id3 have impaired Th17 cell development in mouse models of experimental autoimmune encephalomyelitis and asthma, respectively (26, 28). During viral infection, Th1 cells preferentially express Id2 while Tfh cells express high levels of Id3 (21, 29). Id2-deficiency impairs Th1 differentiation after infection and leads to the generation of effector cells with mixed Th1/Tfh characteristics while lack of Id3 promotes Tfh development following infection and immunization (21, 30).

While mRNA levels are reported as proxy for protein abundance and activity of corresponding proteins; localization, post-translational modifications and programmed destruction of proteins also contribute to regulation of protein concentration within the cell (31, 32). Ubiquitylation is a key post-translational modification that facilitates protein-protein interactions, directs protein localization, instructs protein destruction, and regulates signal transduction, all of which are essential for proper cell function (33, 34). Ubiquitylation is achieved through a regulated, multi-step enzymatic reaction with E3 ubiquitin ligases catalyzing the formation of ubiquitin chains and deubiquitinase (DUB) enzymes countering this by removing ubiquitin modifications in a substrate-specific manner (33-35). Recently, a DUB family member, USP1, was shown to play a major role in promoting Id2 stability in a common bone cancer, osteosarcoma (36). Overexpression of USP1 *in vitro* resulted in increased abundance of non-ubiquitinated Id2. In contrast, USP1 knockdown experiments in osteosarcoma cell lines and primary osteoblasts resulted in Id2 protein destabilization and increased E protein transcriptional activity (36). Further analysis of several primary osteosarcoma tumors also showed a significant increase in USP1 expression that correlated with an increase in Id2 levels (36).

As the exact mechanisms regulating E/Id protein activity that maintain controlled expansion, differentiation and survival of T cells remain unclear, we investigated the role of Usp1 in antiviral T cell responses. Here, we show Usp1 is expressed by effector CD4^+^ and CD8^+^ T cells and interacts with both Id2 and Id3 proteins. Despite this interaction, Usp1 did not appear to be essential for regulation of Id protein abundance and was not required for expansion or differentiation of CD4^+^ or CD8^+^ T cells in primary infection; however, Usp1-deficiency moderately impaired the accumulation of secondary effector populations. Thus, in contrast to its role in osteosarcoma, Usp1 is not necessary for Id2 protein stability in T cells (36).

## Materials and Methods

### Mice and cell lines

All mouse strains were bred and housed in specific pathogen–free conditions in accordance with the Institutional Animal Care and Use Guidelines of the University of California San Diego. Usp1 knockout-first gene-targeted C57BL/6 ES cells were obtained from the KOMP Repository (www.komp.org) and were created at the University of California Davis by the trans-NIH Knock-Out Mouse Project (KOMP) using vector Usp1^tm1a(KOMP)Wtsi^ (37). Usp1-LacZ reporter mice were generated by the Transgenic Mouse and Knock-Out Core at UCSD. Usp1^f/f^ mice were produced by mating the Usp1-LacZ reporter mouse line to FLPo deleter strain (The Jackson Laboratory, stock no 012930). Usp1-floxed mice were subsequently crossed to a CD4-Cre recombinase line (The Jackson Laboratory, stock no 017336) for T cell-specific deletion of Usp1. Id2^fl/fl^ (38), Id3^fl/fl^(39), P14 mice (with transgenic expression of H-2D^b^–restricted TCR specific for LCMV glycoprotein gp33), CD45.1 and CD45.1.2 congenic mice were all bred in house. All mice were fully backcrossed onto a C57BL/6J background. Both male and female mice were used throughout the study, with sex- and age-matched T cell donors and recipients. Mouse A20 cells (ATCC TIB-208) were grown in RPMI-1640 media supplemented with 10% fetal bovine serum and 0.05mM 2-mercaptoethanol.

### T cell activation

Naive CD4^+^ or CD8^+^ T cells were positively selected from single-cell suspensions prepared from spleen and lymph node by mechanical disruption using streptavidin MicroBeads according to manufacturer’s instructions (Miltenyi Biotec). Purified cells were activated in twelve-well plates coated with 0.1-10 μg/mL αCD3 (145–2C11, eBioscience), and 1 μg/mL αCD28 (37.51, eBioscience) for indicated time. To assess proliferation, naive cells were stained with CellTrace Violet according to manufacturer’s protocol (Invitrogen) prior to activation.

### Immunoprecipitation and Western Blot

Proteins were extracted in lysis buffer (1% NP-40, 120 mM NaCl, 50 mM Tris-HCl [pH 7.4], and 1 mM EDTA) containing protease inhibitor cocktail (Sigma). 10ug of protein per sample was resolved on NuPage 4-12% Bis-Tris precast gels in MES buffer (Invitrogen), transferred to 0.45μm PVDF membrane then blocked with 5% BSA in TBS supplemented with 0.1% Tween-20. Id2 (1:1000; 9-2-8, CalBioeagents or D39E8, Cell Signaling), Id3 (1:1000; 6-1, CalBioreagents), Usp1 (1:1000; D37B4, Cell Signaling), Wdr48 (1:1000; NBP1-81404, Novus Biologicals) or β-actin (1:1000; 8H10D10, Cell Signaling) primary antibodies were incubated overnight at 4°C followed by HRP-conjugated secondary antibodies for 1 hour at room temperature (1:10000, Jackson ImmunoResearch). Proteins were visualized with chemiluminescent ECL Prime Western Blotting Detection Reagent (Amersham) or ECL Western Blotting Substrate (Pierce) and imaged on a BioRad ChemiDoc. ImageJ software was used to quantify protein bands. For Immunoprecipitation, 1-1.5 mg of protein lysate was incubated with Id2 (9-2-8, CalBioeagents) or Id3 (6-1, CalBioreagents) antibody overnight at 4°C followed by protein A/G agarose beads (Santa Cruz) for 2-3 hours at 4°C.

### Cell transfer and infections

P14 CD8^+^ T cells congenically distinct for CD45 were adoptively transferred at 5×10^4^ cells per recipient mouse. Mice were then infected with 2×10^5^ PFU LCMV-Armstrong by intraperitoneal injection, 2×10^6^ PFU VV-GP_33_ by intraperitoneal injection or 3×10^4^CFU Lm-GP_33_ by intravenous injection.

### Flow Cytometry and Sorting

Single-cell suspensions were prepared from blood or spleen. The following antibodies were used for surface staining (all from eBioscience unless otherwise stated): CD4 (RM4-5 or GK1.5), CD8 (53-6.7), CD11a (M17/4, Biolegend), CD27 (LG-7F9), CD43 (1B11), CD44 (IM7,eBioscience or MEL-14,Biolegend), CD45.1 (A20-1.7), CD45.2 (104), CD49d (R1-2), CD62L (MEL-14), CD122 (TM-b1), CD127 (A7R34), CXCR3 (CXCR3-173), CXCR5 (SPRCL5, Invitrogen), KLRG1 (2F1), PD1 (J43), and SLAM (TC15-12F12.2, Biolegend). GP_33_ (H-2D^b^/KAVYNFATC, NIH) or GP_276_ (H-2D^b^/SGVENPGGYCL, NIH) tetramers were included with the surface antibody stain. Cells were incubated for 30 min at 4°C in PBS supplemented with 2% bovine growth serum and 0.1% sodium azide. Intracellular staining was performed using the BD Cytofix/Cytoperm Solution Kit (BD Biosciences) and the following antibodies: BCL2 (3F11, BD PharMingen), Eomes (Dan11mag, eBiosciences), GzmB (GB12; Invitrogen), Ki67 (SolA15, eBiosciences), Tbet (eBio4B10, eBiosciences), and TCF1 (C63D9, Cell Signaling). For cytokine staining, splenocytes were incubated for 5 hours at 37°C in RPMI-1640 media containing 10% (v/v) bovine growth serum with 10 nM GP_33-41_ peptide (Anaspec) and Protein Transport Inhibitor (eBioscience) then stained with IFNγ (XMG1.2, eBioscience) and TNFα (MP6-XT22, eBioscience) antibodies. For LacZ reporter detection by flow cytometry, cells were loaded with Fluorescein di(β-D-galactopyranoside) (FDG, Sigma). Single cell suspensions in warm HBSS buffer (1x HBSS supplemented with 2% fetal calf serum and 10mM buffer [pH 7.2]) were mixed in a 1:1 ratio with 2mM FDG diluted in dH_2_O and incubated at 37°C for 1 min. The cell mixture was added to ice cold HBSS buffer and incubated for 1.5 hours on ice before staining with surface antibodies. Stained cells were analyzed using LSRFortessa or LSRFortessa X-20 cytometers (BD) and FlowJo software (TreeStar). All sorting was performed on BD FACSAria or BD FACSAria Fusion instruments.

### Statistics

Two-tailed paired Student’s t test and linear regression analyses were performed using GraphPad Prism software. Microarray data from Immgen (https://www.immgen.org/) was analyzed using GenePattern software.

## Results

### Usp1 is expressed upon T cell activation

Usp1 has recently been shown to play a major role in promoting Id2 stability and subsequent regulation of E protein activity in osteosarcoma (36). Given the necessity of Id2 for T cell survival and differentiation following infection (16, 18, 20-22) and the corresponding increase in Usp1 mRNA upon CD4^+^ and CD8^+^ T cell activation (Fig. 1A), we further investigated the role of Usp1 as a regulator of T cell fate and survival.

**Figure 1.**
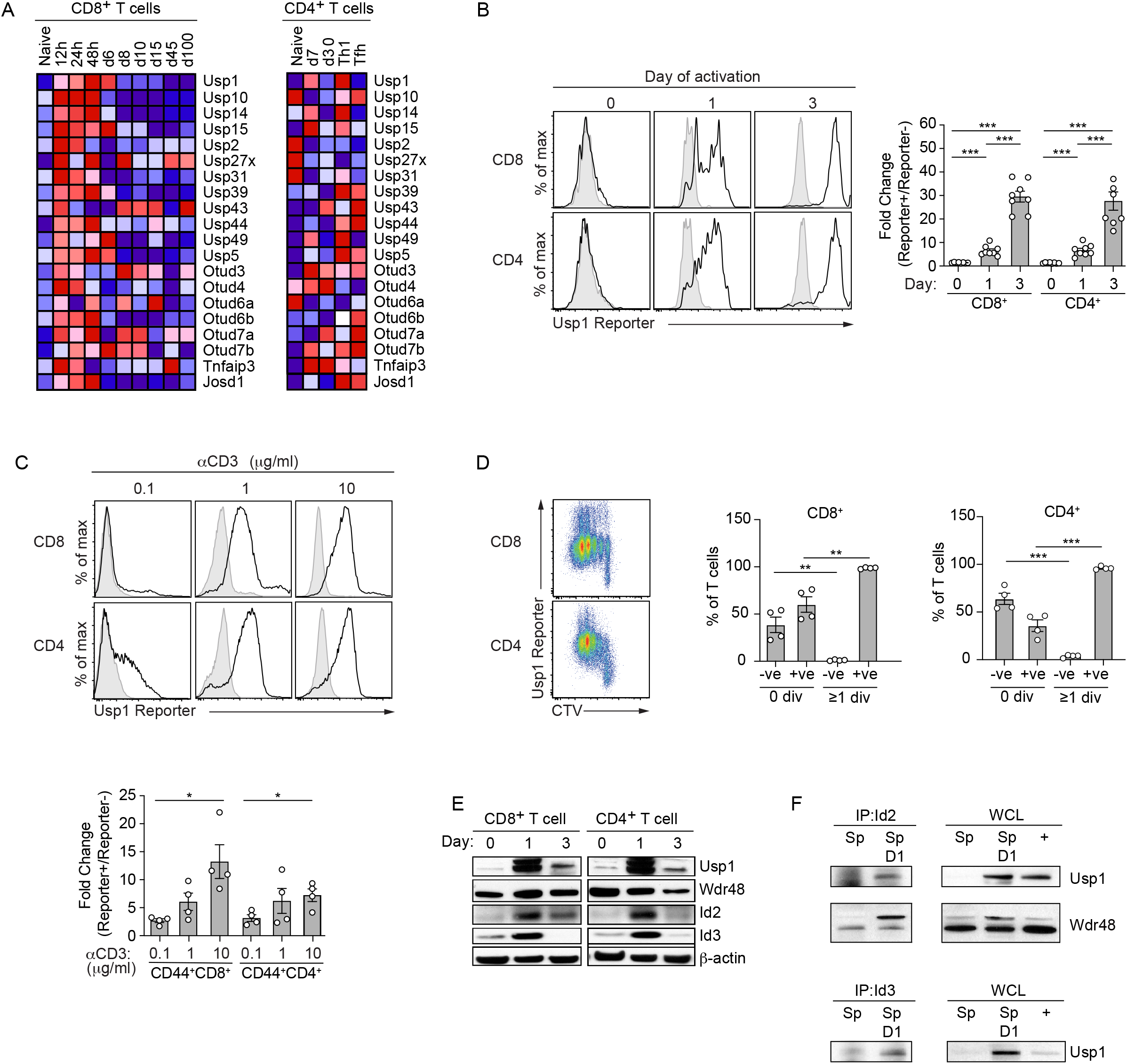
Usp1 is expressed and interacts with Id proteins in activated T cells. **(A)** Heatmap illustrating the relative expression of deubiquitinases differentially expressed in antigen-specific CD8^+^ T cells (left) or CD4^+^ T cells (right) at various time of LCMV infection. **(B-C)** Purified naive wild type (WT) or Usp1^Lacz/+^ CD8^+^ or CD4^+^ T cells were *in vitro* activated with 10μg/ml of αCD3 plus αCD28 for indicted days (B) or for 3 days with indicated concentration of αCD3 plus αCD28 (C) and Usp1-LacZ reporter expression was measured in Usp1^Lacz/+^ (Reporter+; black) and WT (Reporter-; grey) T cells by flow cytometry. Quantification of normalization of Usp1-LacZ levels between Reporter+ and Reporter-controls is shown. **(D)** Purified naive Usp1^Lacz/+^ CD8^+^ or CD4^+^ T cells were labelled with cell trace violet (CTV) then *in vitro* activated with 10μg/ml of αCD3 plus αCD28. After 3 days of activation, the T cells were examined by flow cytometry for Usp1-LacZ reporter expression and CTV dilution (left). The proportion of cells expressing Usp1-LacZ reporter (+ve) or not (-ve) in the 0 or ≥1 division is shown (right). **(E)** Purified naive WT CD8^+^ or CD4^+^ T cells were activated as in (B) and on indicated days protein expression was assessed by Western blot analysis. **(F)** Naive WT splenocytes (Sp) or those activated with 10μg/ml of αCD3 plus αCD28 for 1 day were lysed. Immunoprecipitation with Id2 or Id3 antibody was performed followed by Western blot analysis for Usp1 or Wdr48 (left). Usp1 and Wdr48 protein expression was detected in whole cell lysate (WCL) as a control for total protein levels (right). A20 B cell line (+) was used as a control for Usp1 and Wdr48 expression. β-actin was detected as a loading control. Data are from cumulative (B-D) or one representative (E,F) of 2 independent experiments with n=2-4. Graphs show mean ± SEM; *p < 0.05, ** p<0.01, ***p< 0.001.

Usp1 expression may be induced downstream of the T cell receptor (TCR) to regulate the factors, such as Id proteins, that are necessary for effector T cell activation and cell differentiation. We first sought to define the expression pattern of Usp1 in T cells. Using targeted ‘knockout-first’ ES cells, we generated reporter mice where the *LacZ* gene is driven by the Usp1 promoter (37). A flow cytometry-based assay to detect β-galactosidase (*LacZ*) activity in isolated reporter T cells demonstrated Usp1 expression is increased in CD8^+^ and CD4^+^ T cells over 3 days following TCR engagement (Fig. 1B), and with increasing doses of plate-bound αCD3 (Fig. 1C). As well, Usp1 was expressed prior to the first cell division then thereafter in almost all proliferating CD8^+^ and CD4^+^ T cells (Fig. 1D) consistent with the finding in non-immune cells that Usp1 transcription is regulated in a cell-cycle dependent fashion (40).

We confirmed these results by Western blot and compared the expression of Usp1 and its functional partner Wdr48 to that of Id2 and Id3 as these transcriptional regulators have been shown to interact with Usp1 in other cell types (36) and to be necessary for T cell activation and differentiation into effector and memory subsets (Fig. 1E) (16, 18, 20-23). Usp1 protein levels correlated with the abundance of Id2 and Id3 protein, further supporting a role for Usp1 in stabilizing these important T cell factors. We indeed found a direct association of Usp1 with Id2 and Id3 proteins in activated T cells. Usp1 coimmunoprecipitated with both Id2 and Id3 from T cells that had been activated with plate-bound αCD3 for one day (Fig. 1F). While Usp1 undoubtedly targets additional proteins in the T cell activation and differentiation pathways, these results place Usp1 in a setting where it may contribute to Id protein stability and/or function as it does in osteosarcoma cells (36).

### Antigen-specific T cell express Usp1 following infection

Using our established Usp1-LacZ reporter mice, we next examined Usp1 expression patterns in T cells responding to infection. Usp1-LacZ reporter mice were infected with lymphocytic choriomeningitis virus (LCMV) and peripheral blood CD8^+^ T cells specific for the immunodominant epitope GP_33_ (Fig. 2A,C) or the subdominant epitope GP_276_ (Fig. 2B,C) were assessed for Usp1 reporter activity over the course of infection. At day 7 of infection, the peak of the effector CD8^+^ T cell response, ∼90% of all GP_33_-specific CD8^+^ T cells expressed Usp1. On subsequent days, the proportion of CD8^+^ T cells expressing Usp1 steadily declined with ∼8% of CD8^+^ T cells expressing Usp1 by day 15 of infection and only ∼1% of the memory population (>day 30) remained positive for the Usp1 reporter. Within the effector CD8^+^ T cell population responding to the subdominant LCMV epitope, GP_276_, only ∼56% of total cells expressed Usp1 at day 7 of infection suggesting that the strength of TCR signaling may dictate the level of Usp1 expression. Usp1 expression was also transient in the GP_276_-specific population, and similar to the immunodominant T cell population by day 15 of infection only ∼12% of GP_276_-specific CD8^+^ T cells expressed Usp1.

**Figure 2.**
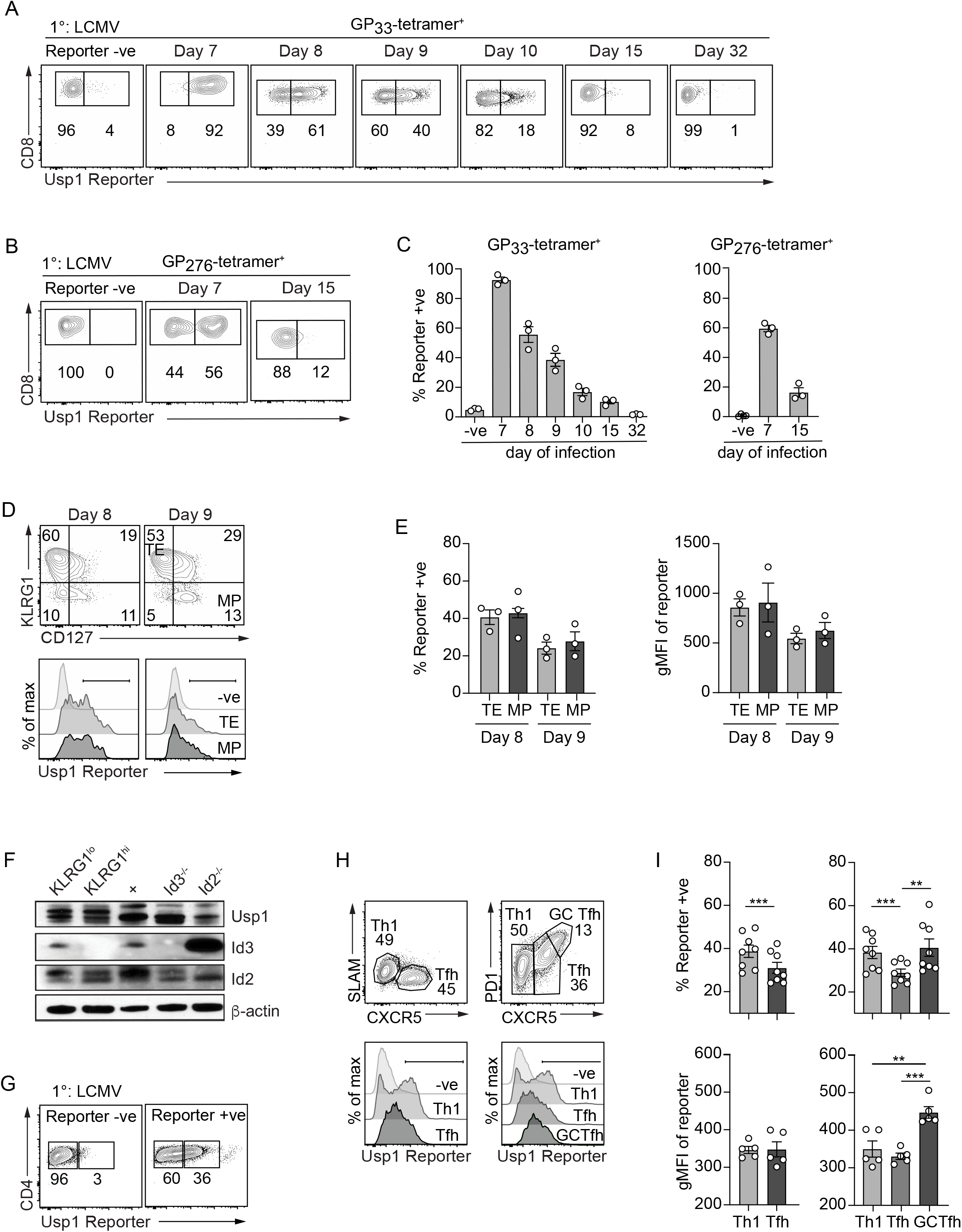
Usp1 is expressed in antigen-specific effector T cells following infection. Usp1^Lacz/+^ reporter or wild type (Reporter -ve) mice were infected with LCMV. The **(A)** GP_33_-specific or **(B)** GP_276_-specific CD8^+^ T cells in the peripheral blood were examined by flow cytometry for Usp1-LacZ reporter expression at indicated time of infection. **(C)** Quantification of the frequency of indicated tetramer^+^ CD8^+^ T cells expressing Usp1-LacZ reporter. **(D)** Terminal effector (TE; KLRG1^+^CD127^-^) and memory precursor (MP; KLRG1^-^CD127^+^) CD8^+^ T cells at day 8 and 9 of infection were examined by flow cytometry for Usp1-LacZ reporter expression. **(E)** Quantification of the frequency of TE or MP cells expressing Usp1-LacZ reporter (left) or the gMFI of Usp1-LacZ reporter expression in the total TE and MP populations (right) is shown. **(F)** On day 10 of infection, KLRG1^hi^ and KLRG1^lo^ CD8^+^ T cells were sort purified from the spleen and Usp1, Id3 and Id2 protein levels were assessed by Western blot analysis. A20 B cells were used as a control for Usp1 expression. Id3^-/-^ and Id2^-/-^ thymocytes were used as a control for Id2 and Id3 expression. β-actin expression was analyzed for a loading control. Total **(G)** or Th1 (SLAM^+^CXCR5^-^ or PD1^-^CXCR5^-^), Tfh (SLAM^-^CXCR5^+^ or PD1^-^CXCR5^+^) and GC-Tfh (PD1^+^CXCR5^+^) subsets **(H)** of CD11a^+^CD49d^+^ antigen-experienced CD4^+^ T cells from the spleen at day 8 of infection were examined by flow cytometry for Usp1-LacZ reporter expression. **(I)** Quantification of the proportion of Reporter+ cells (top) or the gMFI of Usp1-LacZ reporter expression (bottom) in the CD4^+^ T cell subsets is shown. Numbers in plots indicate percent of cells in corresponding gate. Data are from one representative (A-H) or cumulative (I) of 2 independent experiments with n=3-5. Graphs show mean ± SEM; **p < 0.01, ***p < 0.001.

CD8^+^ T cells subsets are differentially reliant on Id protein transcriptional regulation with Id2 promoting and sustaining survival and terminal differentiation of the terminal effector (TE; KLRG1^hi^ CD127^lo^) CD8^+^ T cells (16, 18, 20-22) and Id3 (22, 23) supporting memory precursor (MP; KLRG1^lo^CD127^hi^) effector and memory CD8^+^ T cell populations. While Id3 expression is specific to the MP CD8^+^ T cells, Id2 is expressed in all effector CD8^+^ T cell populations leading us to speculate that subset-specific protein stability and/or function mediated by DUBs might account for the TE subset dependency on Id2. Thus, we next compared Usp1 expression between TE and MP CD8^+^ T cells in the spleen (Fig. 2D). The proportion of Usp1 expressing cells (Fig. 2E, left) and the degree to which Usp1 was expressed (Fig. 2E, right) was comparable between TE and MP CD8^+^ T cells at day 8 and 9 of infection. Western blot analysis of KLRG1^hi^ and KLRG1^lo^ CD8^+^ T cells at day 10 of infection confirmed similar Usp1 expression across effector CD8^+^ T cell subsets (Fig. 2F).

Splenic effector CD4^+^ T cells from Usp1-LacZ reporter mice infected with LCMV-Armstrong were also analyzed. On day 8 of infection, ∼36% of antigen-experienced (CD11a^+^CD49d^+^) CD4^+^ T cells expressed Usp1 (Fig. 2G). We previously reported differential Id2 and Id3 expression among CD4^+^ T cell subsets with Th1 cells exhibiting robust Id2 expression and Tfh and GC-Tfh cells predominantly expressing Id3 (21). We further examined these subsets to see if Usp1 was also differentially expressed (Fig. 2H,I). Th1 cells (CXCR5^-^ SLAM^+^ or CXCR5^-^ PD1^-^) had clear bimodal expression of Usp1; ∼40% of cells within the Th1 population were Usp1 reporter^+^. The Tfh subset (CXCR5^+^SLAM^lo^ or CXCR5^+^PD1^lo^) contained the fewest Usp1 expressing cells (Fig. 2I, top); and the GC-Tfh population showed more homogeneous Usp1 reporter expression than the Th1 subset, with the highest mean fluorescence of the Usp1 reporter (Fig. 2I bottom). Overall, CD4^+^ T cell subsets responding to LCMV infection showed unique patterns of Usp1 expression; however, this expression was not as distinct between the Th1 and Tfh populations as that exhibited by the Id proteins (21).

Our *in vitro* analysis of Usp1 expression indicated Usp1 was expressed in T cells that had divided (Fig.1D). To correlate Usp1 expression to differences in effector T cell expansion, we next examined the population of antigen-specific T cells responding to three distinct pathogens. To track the T cell responses *in vivo*, the Usp1-LacZ reporter mice were crossed to LCMV-GP_33_-specific P14 TCR transgenic mice. Congenically distinct Usp1-LacZ reporter P14 CD8^+^ T cells were transferred into mice that were then infected with LCMV or 2 different recombinant pathogens, *Listeria monocytogenes* bacterial strain (Lm-GP_33_) or recombinant Vaccinia Virus strain (VV-GP_33_). Peripheral blood CD8^+^ T cells compared over the course of infection were examined by flow cytometry for Usp1-LacZ reporter activity (Fig. S1A,B). CD8^+^ T cell population responding to LCMV infection exhibited the earliest Usp1 expression and the greatest proportion of Usp1-expressing cells, with ∼95% at day 6 of infection. Lm-GP_33_ infection induced a peak of Usp1 expression in effector CD8^+^ T cells a day later, at day 7 of infection, with ∼53% of all P14 CD8^+^ T cells expressing Usp1. Finally, the frequency of Usp1 expression was lowest in the CD8^+^ T cell population responding to VV-GP_33_ infection with ∼37% of cells expressing Usp1 at day 8 of infection. Importantly, the degree of expansion of CD8^+^ T cells correlated with the frequency of Usp1 expressing cells (Fig. S1C) indicating that the extent of T cell proliferation following infection likely dictates the extent of Usp1 expression. Regardless of the frequency of Usp1 expressing T cells within the effector T cell population, Usp1 expression was similar between the TE and MP subsets in all infections examined (Fig. S1D,E).

### TE CD8^+^ T cells express increased Usp1 in recall responses

Upon multiple encounters with antigen, the effector CD8^+^ T cell population evolves to include an increased frequency of KLRG1 and Granzyme B expressing T cells as well as acquires a slower ability to attain central memory characteristics and displays stepwise changes in its transcriptional memory program (3, 41-44). Thus, we next examined Usp1 expression in a CD8^+^ T cell population responding to rechallenge. Congenically distinct naive Usp1-LacZ reporter P14 CD8^+^ T cells were transferred into mice that were then infected with LCMV. After 30 days of infection, the mice were rechallenged with Lm-GP_33_ and the antigen-specific CD8^+^ T cells were examined for Usp1 expression over the course of the secondary response. The mice were infected a third time with a VV-GP_33_ infection to assess a tertiary response. The CD8^+^ P14 population in the peripheral blood had the highest frequency of Usp1-expressing cells at day 4 and 5 following both secondary and tertiary infection with a peak of 27% and 23% (Fig. 3A,B), respectively. Interestingly, when the TE and MP were individually assessed for Usp1 expression, on both day 4 and 5 of the secondary or tertiary infection, the TE CD8^+^ T cell subset had a significantly increased frequency of Usp1-expressing cells compared to the MP population on the same day of infection (Fig. 3C,D). To ensure this phenomenon was not simply due to differences in the pathogens used for rechallenge, we transferred congenically distinct Usp1-LacZ reporter P14 CD8^+^ T cells into mice then infected half with LCMV and the other half with Lm-GP_33_. After 30 days, these mice were rechallenged with the opposite pathogen to that used for the primary infection. Regardless of whether LCMV or Lm-GP_33_ was delivered as a secondary infection, the TE subset contained an increased frequency of Usp1-expressing CD8^+^ T cells compared to the MP population (Fig. S2).

**Figure 3.**
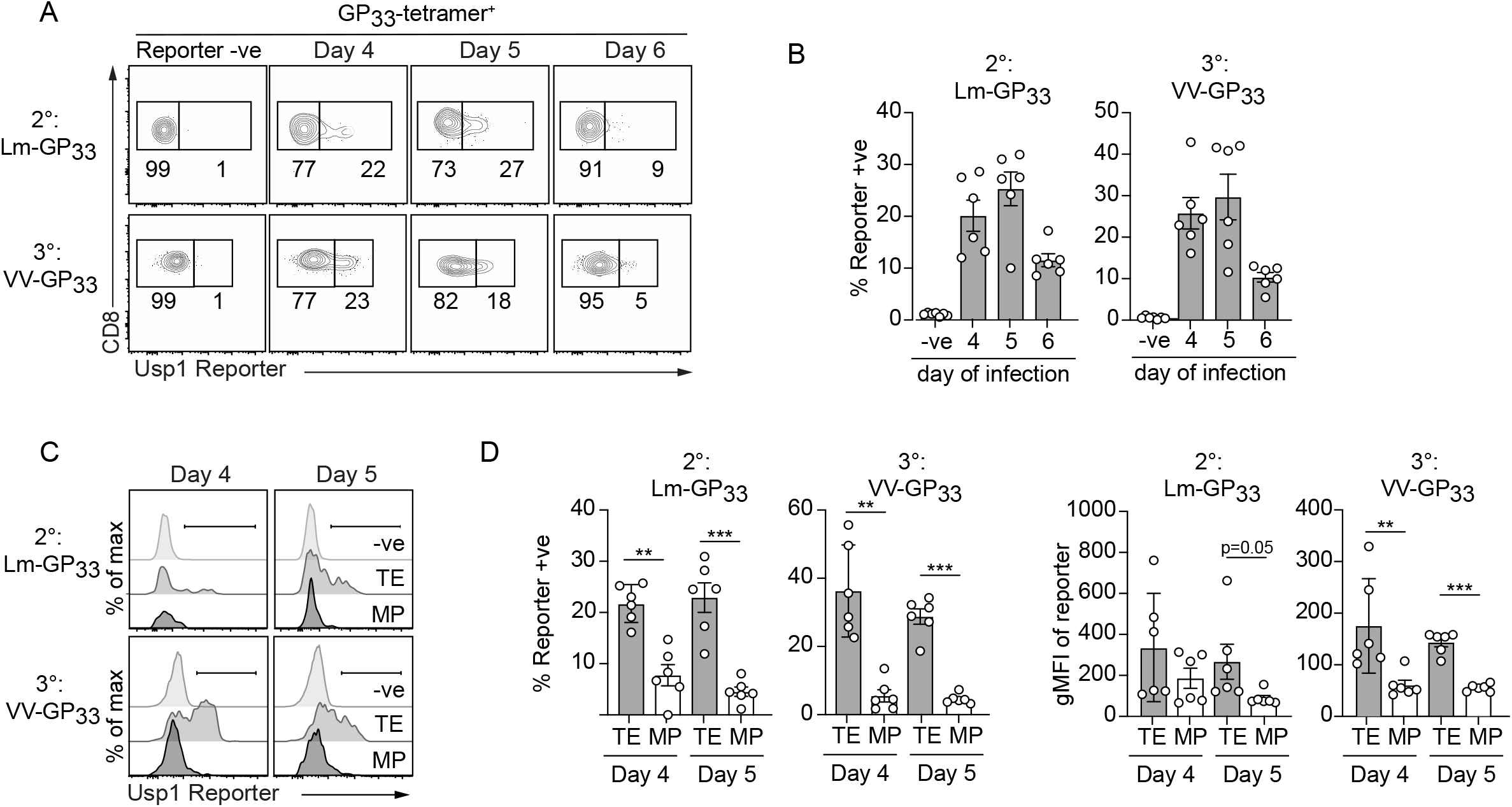
Terminal effector CD8^+^ T cells express increased levels of Usp1 during recall responses. Usp1^Lacz/+^ reporter or wild type (Reporter -ve) mice with resolved primary LCMV infections were rechallenged with a secondary Lm-GP_33_ (2°) and a subsequent tertiary VV-GP_33_ (3°) infection. **(A)** Usp1-LacZ reporter expression in total peripheral blood GP_33_-specific CD8^+^ T cells on day of indicated infection is shown. **(B)** Quantification of the frequency of indicated tetramer^+^ CD8^+^ T cells expressing Usp1-LacZ reporter. **(C)** TE (KLRG1^+^CD127^-^) and MP (KLRG1^-^ CD127^+^) effector CD8^+^ T cell populations on the indicated day of infection were examined by flow cytometry for Usp1-LacZ reporter expression. **(D)** Quantification of the frequency of Reporter^+^ cells (left) or the gMFI of Usp1-Reporter (right) in the indicated subset is shown. Numbers in plots indicate percent of cells in corresponding gate. Data are cumulative of 2 independent experiments with n=3. Graphs show mean ± SEM; **p < 0.01, ***p < 0.001.

### Usp1-deficient CD8^+^ T cells fail to accumulate during rechallenge

We next assessed the role of Usp1 in survival and differentiation of T cells responding to infection. To achieve this, we generated Usp1 conditional knockout (KO) mice by crossing our Usp1-LacZ reporter mouse line to transgenic *FLP* mice. Flp recombinase converts the Usp1 knockout-first allele to a Usp1 conditional allele that is floxed with loxP sites (37). The Usp1-floxed mice were then crossed to a CD4-Cre recombinase line to achieve T cell-specific deletion of Usp1. Finally, these mice were crossed to the P14 TCR transgenic line so we could track antigen-specific T cells *in vivo*. Congenically distinct Usp1^f/f^-CD4Cre^+^ (Usp1^KO^) and wildtype littermate (Usp1^WT^) P14 CD8^+^ T cells were mixed 1:1 and transferred into recipient mice that were then infected with LCMV. The frequency of Usp1^KO^ P14 CD8^+^ T cells remained equivalent or moderately increased compared to Usp1^WT^ cells in the peripheral blood through the effector (day 5-8) phase of infection. The Usp1^KO^ P14 CD8^+^ T cells persisted into the memory (∼day 60-74) phase of infection at a marginally reduced frequency compared to WT controls (Fig. 4A,B). Usp1 deficiency had minimal impact on proliferation and survival of effector CD8^+^ T cells at day 5 and 7 of primary infection as measured by Ki67 and BCL2 expression, respectively (Fig. 4C). Furthermore, the splenic Usp1^WT^ and Usp1^KO^ antigen-specific effector T cell populations were composed of similar frequencies of TE and MP subsets at day 5 and 7 of infection (Fig. 4D) and expressed comparable levels of T cell activation markers (CD44, PD1), T cell subset defining surface markers (CD27, CD43, CXCR31, and CD122), and transcription factors (TCF1, EOMES, and TBET) (Fig. 4E). As Usp1 has been reported to play a role in DNA damage repair (40, 45), we next assessed the impact of Usp1-deficiency on the CD62L^hi^ MP effector population reported to have heightened DNA damage sensing and repair ability (46). We observed a small, statistically significant increase in this population perhaps suggesting that a reduction of DNA damage repair mechanisms allows for increased cell expansion (Fig. 4F). Finally, effector T cells lacking Usp1 secreted equivalent IFNγ and TNFα following *ex vivo* GP_33_ stimulation and showed elevated expression of Granzyme B compared to Usp1^WT^ P14 CD8^+^ T cells (Fig. 4G,H).

**Figure 4.**
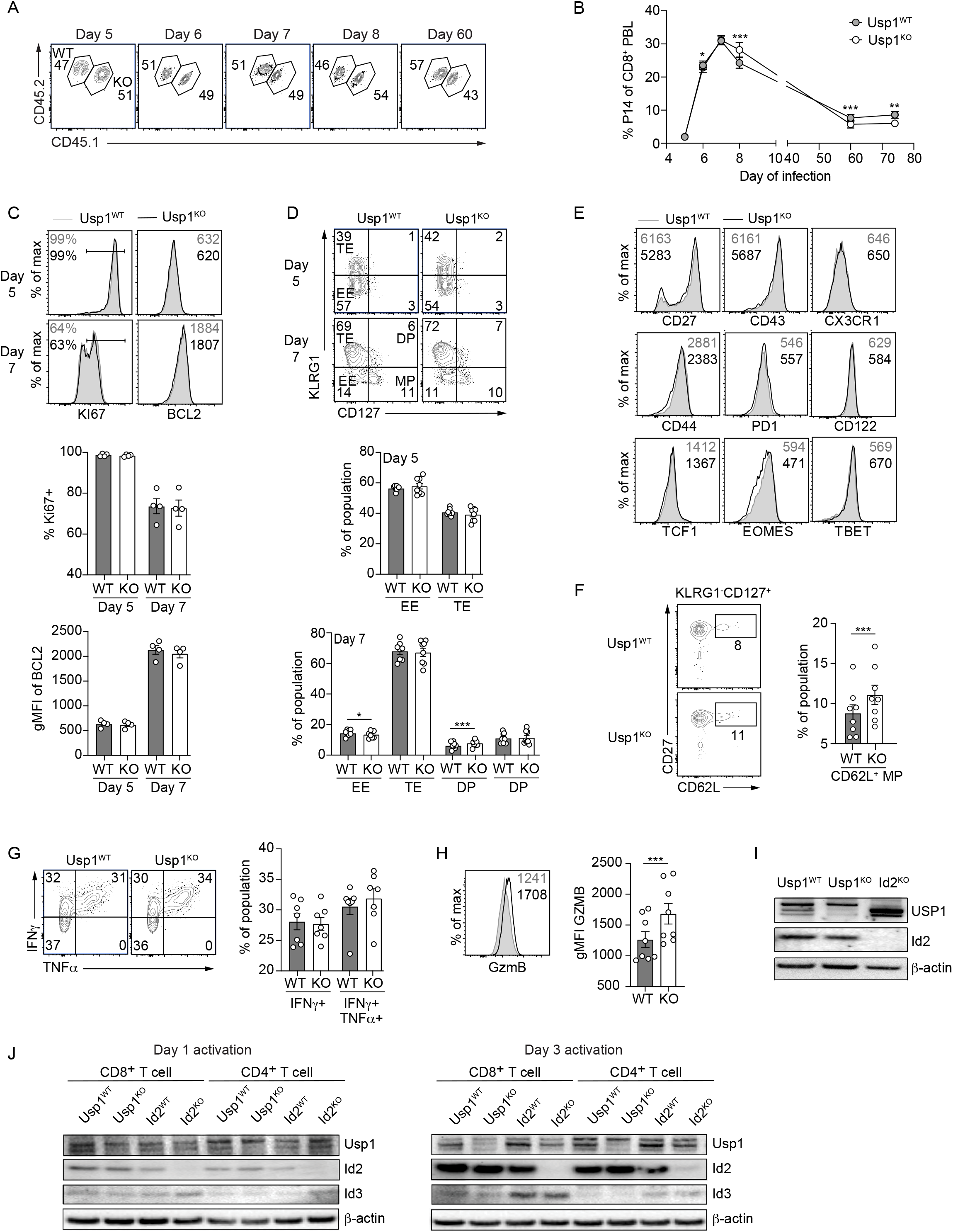
Usp1-deficient CD8^+^ T cells competently respond to primary infection. CD45.1 host mice were given a cotransfer of Usp1^WT^ (CD45.2) and Usp1^KO^ (CD45.1.2) P14 CD8^+^ T cells then subsequently infected with LCMV. **(A)** Frequency of Usp1^WT^ and Usp1^KO^ cells among total P14 CD8^+^ T cells in peripheral blood was assessed by flow cytometry. **(B)** Quantification of the frequency of Usp1^WT^ and Usp1^KO^ P14 CD8^+^ T cells over the course of primary infection is shown. **(C)** Flow cytometry analysis of Ki67 and BCL2 expression in total Usp1^WT^ (grey) and Usp1^KO^ (white) P14 CD8^+^ T cells in the spleen on day 5 and 7 of infection (top). Frequency of Ki67^+^ P14 CD8^+^ T cells and the gMFI of BCL2 expression is quantified (bottom). **(D)** Expression of KLRG1 and CD127 on Usp1^WT^ and Usp1^KO^ P14 CD8^+^ T cells from spleen on indicated day of infection (top). Quantification of the frequency of EE (KLRG1^-^CD127^-^), TE (KLRG1^+^CD127^-^), DP (KLRG1^+^CD127^+^) and MP (KLRG1^-^CD127^+^) cells within the total P14 CD8^+^ T cell population (bottom). **(E)** Expression of indicated surface protein (top, middle) or transcriptional regulator (bottom) in the total Usp1^WT^ (grey) and Usp1^KO^ (white) P14 CD8^+^ T cell population in the spleen on day 5 of infection is shown. **(F)** The expression of CD62L and CD27 among the MP (KLRG1^-^ CD127^+^) population (left). The frequency of the CD62L^+^ subset within the MP (KLRG1^-^CD127^+^) population is shown in spleen on day 7 of infection. **(G)** Total Usp1^WT^ and Usp1^KO^ P14 CD8^+^ T cells from the spleen on day 5 of infection were assessed for IFNγ and TNFα production following *ex vivo* stimulation with GP_33_ peptide (left). Frequency of IFNγ - or IFNγTNFα -producing P14 CD8^+^ T cells is shown (right). **(H)** Granzyme B expression in total Usp1^WT^ (grey) and Usp1^KO^ (white) P14 CD8^+^ T cells from the spleen on day 7 of infection (left). Quantification of the gMFI of granzyme B expression is shown (right). **(I)** Purified naive Usp1^WT^, Usp1^KO^, Id2^WT^ and Id2^KO^ CD8^+^ and CD4^+^ T cells were activated with 10μg/ml of αCD3 plus αCD28 for indicted days or **(J)** total Usp1^WT^ and Usp1^KO^ P14 CD8^+^ T cells were sorted from the spleen at day 7 of LCMV infection. Usp1, Id2 or Id3 protein expression was detected by Western blot. β-actin was detected as a loading control. Thymocytes from Id2^KO^ mice were used as a control for Id2 expression (J). Numbers in graphs indicate percent of cells in corresponding gate (A, C left, D, F, G) or gMFI (C right, E, H) of population. Data are representative (A, C top, D top, E, F left, G left, H left, I, J) or cumulative (B, C bottom, D bottom, F right, G right, H right) of 2 independent experiments with n=4. Graphs show mean ± SEM; **p < 0.01, ***p < 0.001.

While we see the reporter expressed, we do not see an essential cell intrinsic role for Usp1 that correlates with the dependency on Id proteins; thus, we looked to see if Id2 or Id3 protein levels in T cells were altered with Usp1 deficiency. Naive CD8^+^ and CD4^+^ T cells lacking Usp1 were activated *in vitro* with αCD3 and αCD28 then Id2 and Id3 protein was detected by Western blot. Following activation, Id2 and Id3 protein levels in Usp1^KO^ CD4^+^ and CD8^+^ T cells were comparable to corresponding wild type cells (Fig. 4I). Furthermore, Usp1^WT^ and Usp1^KO^ CD8^+^ P14 cells at day 7 of LCMV infection also expressed equivalent levels of Id2 protein (Figure 4J). However, when we assessed Id2 and Id3 protein levels in Usp1-deficient thymocytes, we detected a slight decrease in Id2 protein expression (Fig. S3). Therefore, while Usp1 expression was dynamically regulated in activated T cells, loss of Usp1 alone did not appear to be essential for Id protein stability in effector T cells, in contrast to thymocytes and osteosarcoma cells (36).

As noted above, Usp1-deficient P14 CD8^+^ T cells declined in frequency compared to WT cells at late memory timepoints (Fig. 4A,B). In particular, a decreased proportion of Usp1^KO^ T_CM_ (CD62L^hi^CD127^hi^) was observed despite an increased frequency CD62L^+^ MP cells observed earlier in infection (Fig. 5A). Furthermore, Usp1 expression appeared more dynamic across the secondary and tertiary effector CD8^+^ T cell subsets as compared to the primary effector populations (Fig. 3); thus, we next evaluated the recall response of Usp1-deficient memory CD8^+^ T cells. Mice receiving a mixed transfer of Usp1^WT^ and Usp1^KO^ P14 CD8^+^ T cells were infected with LCMV then 30 days after infection were rechallenged with Lm-GP_33_. Following the kinetics of the CD8^+^ T cell secondary response in the blood, we found the Usp1^KO^ CD8^+^ T cells showed a modest accumulation defect following reinfection and were maintained at a reduced frequency compared to the Usp1^WT^ cells beyond the contraction phase (Fig. 5B,C). The Usp1^KO^ secondary effector CD8^+^ T cell population contained a slight but significantly increased frequency of KLRG1^hi^ cells (Fig. 5D) that was reflected in a somewhat increased proportion of DP (KLRG1^hi^CD127^hi^) Usp1^KO^ effector T cells compared to the Usp1^WT^ control at day 4 and 5 of secondary infection (Fig. 5E). Consistent with a more differentiated phenotype, Usp1^KO^ CD8^+^ T cells had decreased CD27 and increased CX3CR1 levels over Usp1^WT^ cells (Fig. 5F). Thus, although modest, Usp1-deficiency impacted the secondary effector T cells more so than the effector T cells of the primary response suggesting stage-specific regulation of T cell populations.

**Figure 5.**
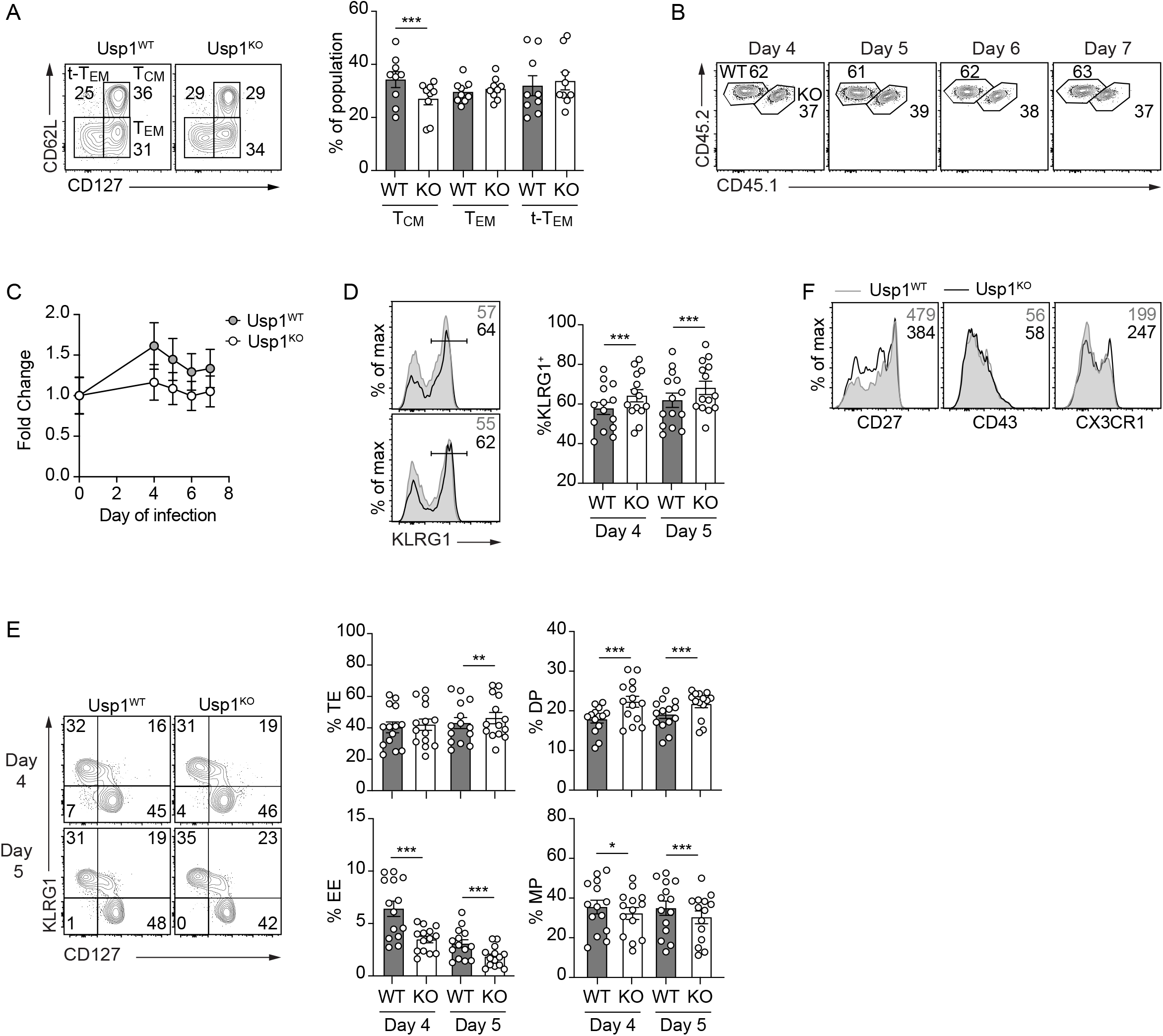
Usp1-deficient CD8^+^ T cells have an impaired secondary response. **(A)** Expression of CD62L and CD127 on Usp1^WT^ and Usp1^KO^ P14 CD8^+^ T cells in the blood at >60 days of LCMV was measured by flow cytometry (left). Quantification of the frequency of T_CM_ (CD62L^+^CD127^+^), T_EM_ (CD62L^-^CD127^+^), t-T_EM_ (CD62L^-^CD127^-^) subsets of total P14 CD8^+^ T cells in the blood (right). **(B-F)** Recipient mice were rechallenged with Lm-GP_33_ at >30 days of the primary LCMV infection. **(B)** Frequency of Usp1^WT^ and Usp1^KO^ cells among total P14 CD8^+^ T cells in peripheral blood was assessed by flow cytometry. **(C)** Quantification of the fold change in frequency of Usp1^WT^ and Usp1^KO^ P14 CD8^+^ T cells at indicated day of infection compared to pre-rechallenge levels is shown. **(D)** KLRG1 expression on Usp1^WT^ (grey) and Usp1^KO^ (white) P14 CD8^+^ T cells from the blood on indicated day of expression (left). Quantification of the frequency of KLRG1^+^ cells is shown (right). **(E)** Expression of KLRG1 and CD127 on Usp1^WT^ and Usp1^KO^ P14 CD8^+^ T cells of the blood on indicated day of infection (left). Quantification of the frequency of EE (KLRG1^-^CD127^-^), TE (KLRG1^+^CD127^-^), DP (KLRG1^+^CD127^+^) and MP (KLRG1^-^CD127^+^) cells within the total P14 CD8^+^ T cell population (right). **(F)** Expression of indicated surface protein in the total Usp1^WT^ (grey) and Usp1^KO^ (white) P14 CD8^+^ T cell population in the blood at day 5 of secondary infection is shown. Numbers in graphs indicate percent of cells in corresponding gate (A,B,D,E) or gMFI (F). Data are representative of 2 (A) or 3 (B-E) independent experiments with n=3-6. Graphs show mean ± SEM; *p< 0.05, **p < 0.01, ***p < 0.001.

## Discussion

E and Id protein are key transcriptional regulators driving the development, differentiation and maintenance of T cell populations (47), and thus their expression and function must be precisely modulated. Here, we investigated the regulation of Id2 and Id3 protein stability by the DUB enzyme, Usp1. Usp1 expression was upregulated in activated T cells *in vitro* and in effector CD4^+^ and CD8^+^ T cell populations following infection *in vivo* with the degree of Usp1 expression correlating with the size of the effector CD8^+^ T cell population. Despite its expression pattern, Usp1-deficiency did not impact effector CD8^+^ T cells proliferation and survival and Usp1^KO^ CD8^+^ T cells accumulated to levels comparable to the WT population following primary infection. Differentiation of primary effector CD8^+^ T cells was also unaffected by lack of Usp1 but a gradual loss of memory T cells did occur over time. Furthermore, the subsequent Usp1-deficient secondary effector T cell population had a modest defect in accumulation and exhibited a slightly more differentiated phenotype. While Usp1 associated with both Id2 and Id3 in activated T cells, the Id protein levels in *in vitro* activated T cells and effector CD8^+^ T cells during primary expansion were unaffected by the loss of Usp1. However, we did observe a reduction in Id2 protein levels for Usp1^KO^ thymic T cells as reported for osteosarcomas (36). Taken together, the role of Usp1 in modulating Id protein abundance appears dependent on cell-stage and cell-type, highlighting that additional mechanisms beyond Usp1-mediated deubiquitylation must be employed in effector T cell populations to regulate Id protein turnover.

The fact that T cells within the secondary effector population rely on Usp1 expression to a greater degree than those in the primary effector population may suggest that other DUBs compensate for the absence of Usp1 in the primary response more so than in subsequent responses. Differential regulation of hundreds of genes after repetitive antigen stimulation of CD8^+^ T cells has been reported (44). Id2 expression is upregulated upon each antigen encounter while several DUBs including *Usp36, Usp48, Usp53* and *Stambpl1* decrease in expression with repeated antigen stimulation (44). Increased Id2 protein abundance in secondary effector T cell populations might necessitate increased reliance on deubiquitinase activity while reduced expression of compensating DUB family members may make these T cells more dependent on Usp1. Thus, differential reliance of Usp1 could be explained by dynamic expression of compensatory DUBs across memory populations. Alternatively, Usp1 is also known to function as an important regulator of DNA damage repair processes (40, 45, 48). T cell populations undergoing multiple rounds of proliferation would be subjected to DNA damage and would require mechanisms to maintain genomic integrity that may vary across T cell populations (46). Secondary effector and memory populations may more readily employ Usp1-dependent DNA damage repair mechanisms.

Following activation, dramatic changes in gene expression occur within a T cell to support proliferation, differentiation, effector function and subsequent memory formation. Many studies have contributed to detailing the transcriptional networks that regulate the differentiation of effector and memory T cell population (11, 12, 14, 15). However, despite being essential for defining cell activity and identity, how the transcriptome relates to actual protein abundance or function in T cell populations remains understudied. Studies relating mRNA and protein abundance in T cell populations have been limited by cell numbers and lack of reagents including antibodies specific for E and Id proteins. Importantly, our data here demonstrate Id2 and Id3 protein levels mirror the kinetics of mRNA expression previously described for CD8^+^ T cells (22).

Protein translation is dynamically regulated in CD8^+^ T cell populations responding to acute LCMV infection and tumors (49, 50). A recent study comparing naive and activated human T cells (51) suggest naive T cells exist in a state of preparedness by maintaining a high protein turnover, a large set of idling ribosomes and a pool of repressed messenger RNAs, including those coding for glycolytic enzymes, that can be rapidly engaged in translation. As well, nondegradative and degradative ubiquitylation events have been shown to contribute to various aspects of TCR, costimulatory and coinhibitory, and cytokine signaling that are important for regulating T cell activation, expansion, differentiation and persistence (52). Altogether, this points to the importance of regulation of T cell activation and differentiation at the translation and protein level. Here we examined a very specific function of Usp1 in regulating Id2 protein levels in differentiating T cells, as demonstrated to occur in other cell types (36). While Usp1 appears to play a minor role in antiviral CD8^+^ T cells responses, the notion that fate decision of differentiating T cells is controlled by the rate and quality of protein translation or turnover warrants further study.

## Acknowledgements

We thank Kevin Brannen and Brandon Chechile for their assistance with cell sorting. We also thank Dr. Nicole Scharping and Amir Ferry for critical review of this manuscript. The ES cells used to generate the mouse strain for this research project were created using a vector generated by the trans-NIH Knock-Out Mouse Project (KOMP) and were obtained from the KOMP repository (www.komp.org).

## Grant Support

This study was supported by the National Institutes of Health (NIH) R01 grants (AI067545, AI072117) and the UCSD Tata Chancellor’s Professorship to A.W. Goldrath, the NIH Ruth L. Kirschstein NRSA F31 Predoctoral Fellowship (1F31AG043222-01A1) and the TATA Institute for Genomics and Society Fellowship to L.A. Shaw, and the Leukemia and Lymphoma Society Career Development Award to K.D. Omilusik. NIH grants to Velocigene at Regeneron Inc (U01HG004085) and the CSD Consortium (U01HG004080) funded the generation of gene-targeted vectors and ES cells for over 8500 genes in the KOMP Program and archived and distributed by the KOMP Repository at UC Davis and CHORI (U42RR024244).

## Notes

### Competing Interest Statement

The authors have declared no competing interest.

