## Supplemental Material for "Ubiquitin specific protease 1 expression and function in T cell immunity"

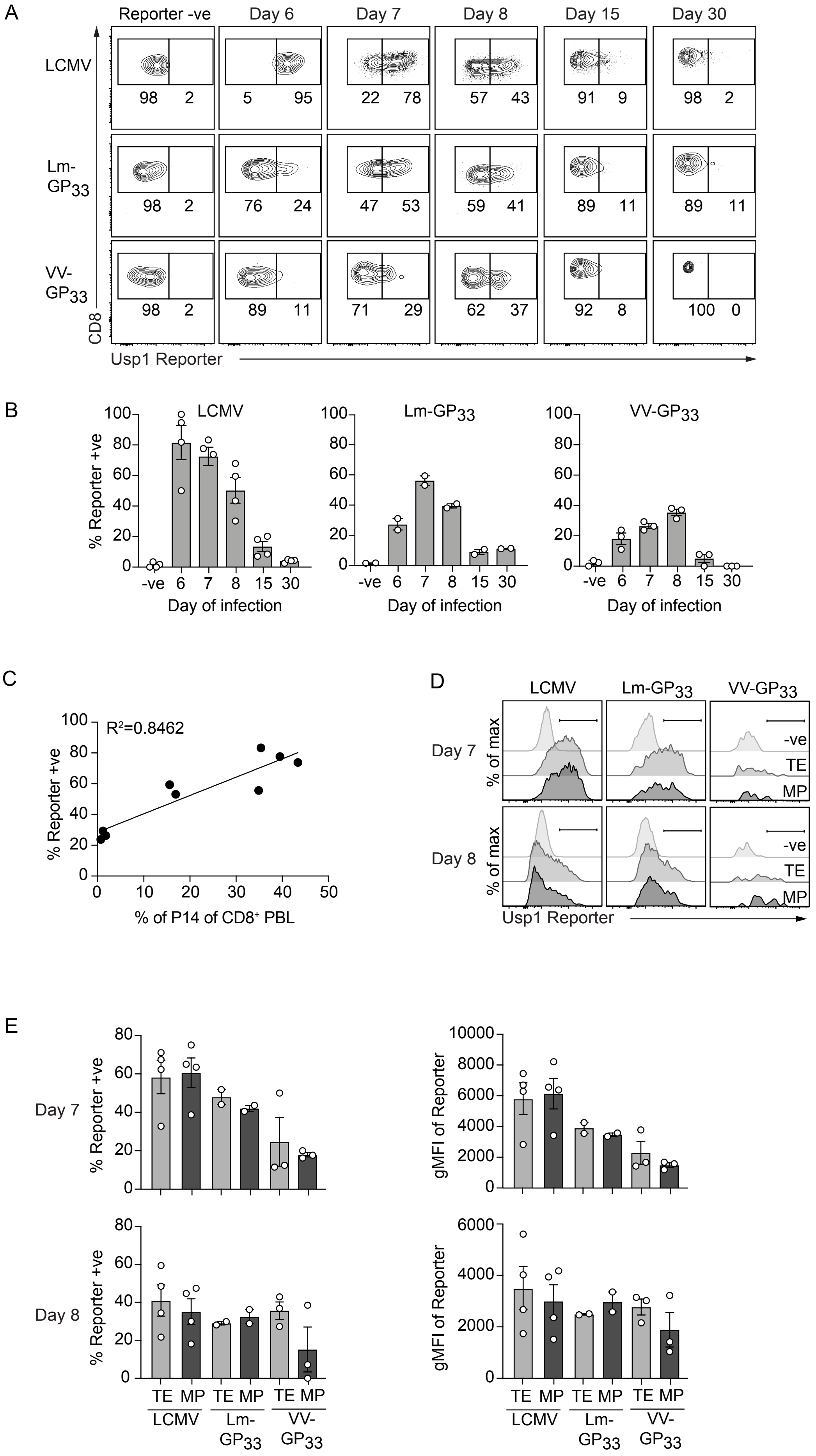
A
Reporter -ve
Day 6
Day 7
Day 8
Day 15
Day 30
LCMV
98 2
5 95
22 78
57 43
91 9
98 2
Lm-GP33
98 2
76 24
47 53
59 41
89 11
89 11
VV-GP33
98 2
89 11
71 29
62 37
92 8
100 0
CD8
Usp1 Reporter
B
LCMV
Lm-GP33
VV-GP33
% Reporter +ve
100
80
60
40
20
0
-ve 6 7 8 15 30
Day of infection
C
R²=0.8462
% Reporter +ve
100
80
60
40
20
0
0 10 20 30 40 50
% of P14 of CD8⁺ PBL
D
LCMV
Lm-GP33
VV-GP33
Day 7
Day 8
% of max
-ve
TE
MP
Usp1 Reporter
E
Day 7
Day 8
% Reporter +ve
80
60
40
20
0
gMFI of Reporter
10000
8000
6000
4000
2000
0
TE MP TE MP TE MP
LCMV
Lm-GP33
VV-GP33

**Supplemental Figure 1. Usp1 expression correlates with degree of effector CD8$^+$ T cell population expansion.** Usp1$^{LacZ/+}$ or wild type (Reporter -ve) P14 CD8$^+$ T cells were transferred to congenically distinct recipients that were subsequently infected with LCMV, Lm-GP$_{33}$ or VV-GP$_{33}$. **(A)** Usp1-LacZ reporter expression was examined by flow cytometry in P14 CD8$^+$ T cells from the blood at indicated days of infection. Quantification of the **(B)** proportion of Reporter$^+$ P14 CD8$^+$ T cells is shown. **(C)** The correlation of frequency P14 CD8$^+$ T cells of the total CD8$^+$ peripheral blood lymphocyte population and the proportion of total P14 CD8$^+$ T cells expressing Usp1-LacZ reporter at day 7 of infection is graphed. **(D)** Terminal effector (TE; KLRG1$^+$CD127$^-$) and memory precursor (MP; KLRG1$^-$CD127$^+$) P14 CD8$^+$ T cells in the blood at day 7 (top) and 8 (bottom) of infection were examined by flow cytometry for Usp1-LacZ reporter expression. Gate for Reporter +ve population is indicated. **(E)** Quantification of the frequency of TE or MP cells expressing Usp1-LacZ reporter (left) or the gMFI of Usp1-LacZ reporter (right) in total P14 CD8$^+$ T cells is shown. Data are cumulative of 1 independent experiments with n=2-4.

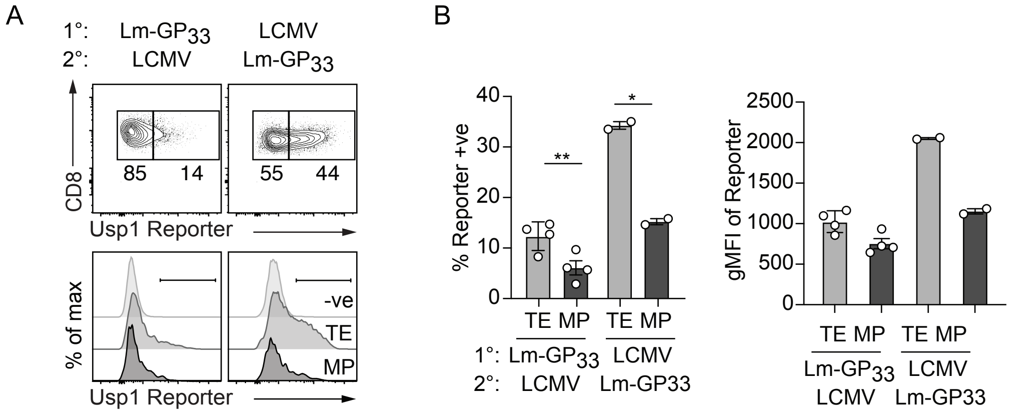


**Supplemental Figure 2. Usp1 expression is increased in TE CD8⁺ T cells regardless of secondary infection.** Usp1$^{LacZ/+}$ P14 T cells were transferred to congenically distinct recipients that were subsequently infected with LCMV or Lm-$GP_{33}$. At >30 days of infection, mice were given the alternate secondary infection as indicated. **(A)** Usp1-LacZ reporter expression in total (top) or in TE and MP (bottom) effector P14 cell populations in the spleen on day 6 of secondary infection is shown. **(B)** Quantification of the frequency of Reporter$^+$ P14 CD8$^+$ T cells (left) or the gMFI of Usp1-LacZ (right) in indicated population is shown. Numbers in plots indicate percent of cells in corresponding gate. Data are cumulative of 1 independent experiments with n=2-4. Graphs show mean ± SEM; $^*p < 0.05$, $^{**}p < 0.01$.

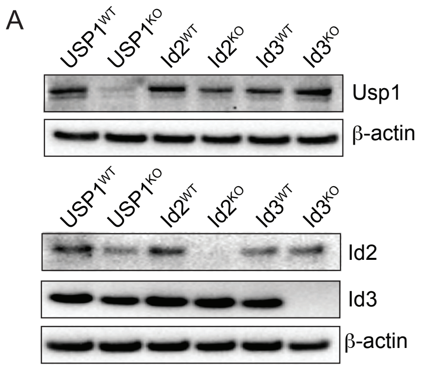


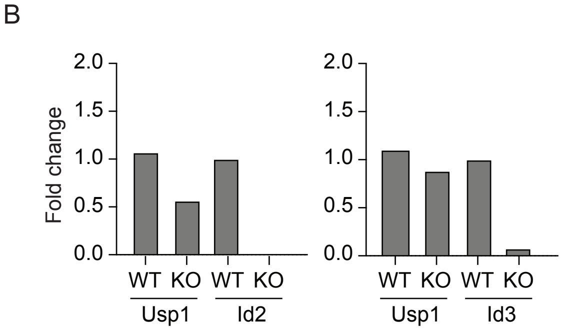


**Supplemental Figure 3 Usp1-deficient thymocytes have reduced Id2 protein levels.**
**(A)** Total $Usp1^{WT}$ and $Usp1^{KO}$, $Id2^{WT}$ and $Id2^{KO}$ or $Id3^{WT}$ and $Id3^{KO}$ thymocytes were examined for Usp1, Id2 or Id3 protein expression by Western blot. β-actin was detected as a loading control. **(B)** Quantification of total Id2 or Id3 protein relative to β-actin loading control normalized to $Id2^{WT}$ or $Id3^{WT}$ sample.
